# *climwin*: An R Toolbox for Climate Window Analysis

**DOI:** 10.1101/069427

**Authors:** Liam D. Bailey, Martijn van de Pol

**Affiliations:** Department of Evolution, Ecology and Genetics, Research School of Biology, The Australian National University, Canberra, Australia; Department of Animal Ecology, Netherlands Institute of Ecology (NIOO-KNAW), Wageningen, The Netherlands

## Abstract

When studying the impacts of climate change, there is a tendency to select climate data from a small set of arbitrary time periods or climate windows (e.g., spring temperature). However, these arbitrary windows may not encompass the strongest periods of climatic sensitivity and may lead to erroneous biological interpretations. Therefore, there is a need to consider a wider range of climate windows to better predict the impacts of future climate change. We introduce the R package ***climwin*** that provides a number of methods to test the effect of different climate windows on a chosen response variable and compare these windows to identify potential climate signals. ***climwin*** extracts the relevant data for each possible climate window and uses this data to fit a statistical model, the structure of which is chosen by the user. Models are then compared using an information criteria approach. This allows users to determine how well each window explains variation in the response variable and compare model support between windows. ***climwin*** also contains methods to detect type I and II errors, which are often a problem with this type of exploratory analysis. This article presents the statistical framework and technical details behind the ***climwin*** package and demonstrates the applicability of the method with a number of worked examples.

## Introduction

With the growing importance of climate change there are an increasing number of studies seeking to understand the impact of climate on biological systems (e.g., [1–5]). However, in many study systems the impacts of climate are likely to be different at different times of the year (e.g., [4–6]), making it necessary for researchers to subset their climate data to encompass a particular period of interest, here termed the climate window (e.g., spring temperature, winter precipitation). However, this subsetting decision is often made with little *a priori* knowledge on the relationship between climate and the biological response, leading to the arbitrary selection of one, or few, climate windows [7].

The use of a limited number of arbitrarily selected climate windows hinders our ability to make meaningful biological conclusions. If a trait, such as body mass or offspring number, displays no response to an arbitrary climate window we cannot determine if this is evidence of climatic insensitivity in our response variable or if the choice of climate window is flawed. Even where we detect a relationship between climate and our response, we cannot know whether there may be another point in time at which climate has a much stronger and more biologically meaningful impact. With flawed conclusions there is a potential to overlook key periods of biological importance, leading us to focus limited management and conservation resources in the wrong areas.

To overcome these issues, there is a need to test a greater number of climate windows with fewer *a priori* assumptions. One solution is the use of a sliding window approach [5, 8–11], where one varies (or slides) the start and end time of a climate window to compare multiple possible windows and select a best window (Fig 1). However, as these analyses are often done manually, comparison of a large number of climate windows can be cumbersome and time consuming. Additionally, there is currently no standardised method for testing or comparing climate windows, and we have no knowledge on the performance of sliding window approaches, including the possibility for false positives and false negatives (type I and II errors); precision and bias of parameter estimates and model statistics (e.g., R^2^); and how these errors and biases might depend on sample size and climate signal strength. There is a need for a standardised and automated approach that can help streamline these frequently performed analyses and make the testing and comparison of multiple climate windows easy and accessible to the general scientific community. The package ***climwin***, built in R, creates a best practice method for this process.

**Fig 1.**
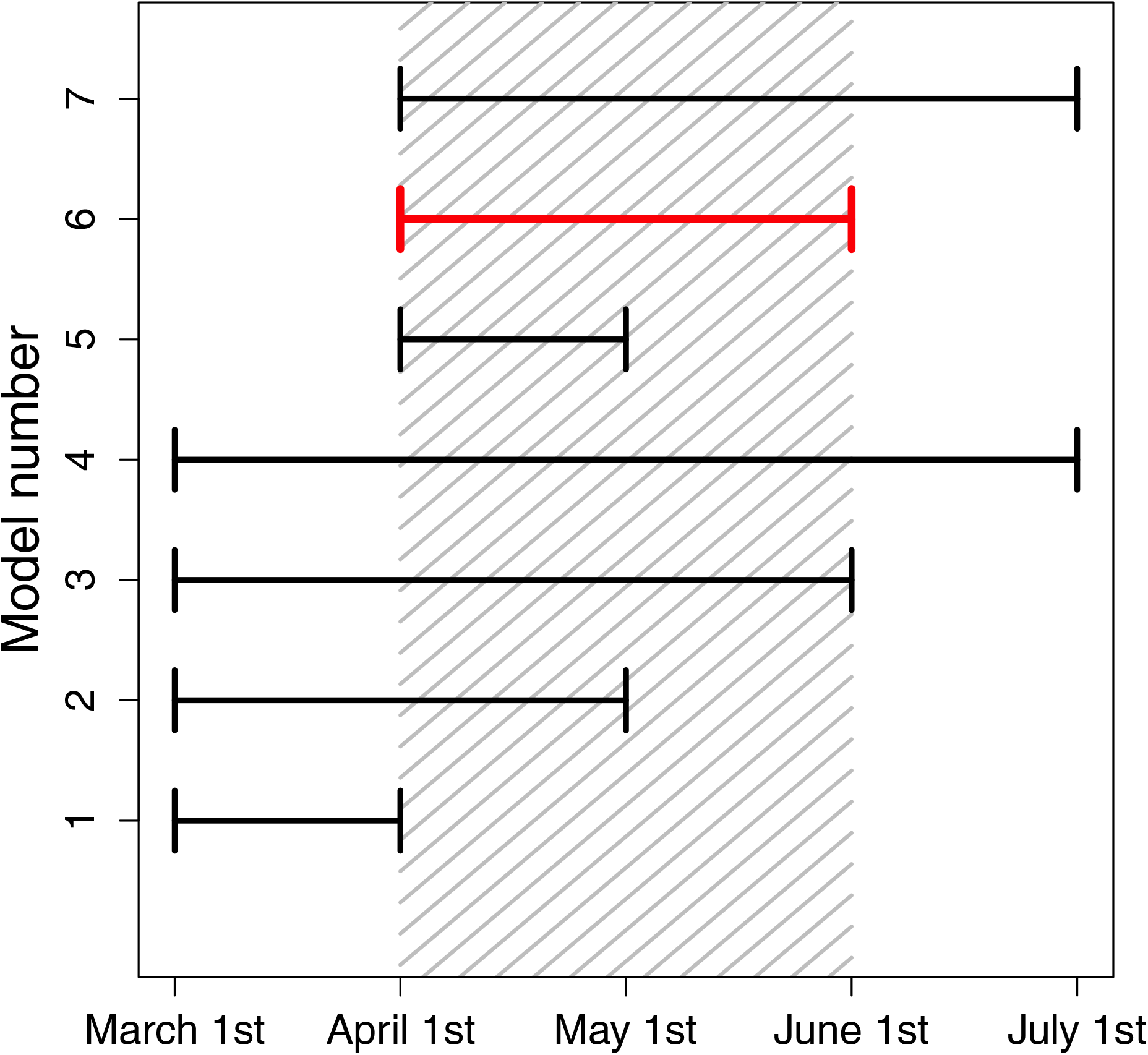
Illustration of a sliding window approach. Shaded region represents a climate signal (April 1^st^ - June 1^st^), where a climatic predictor has the strongest impact on the biological response. Each line represents a tested climate window. The start and end time of windows is varied until we identify the best window (in red). This figure demonstrates a sliding window analysis conducted at a monthly resolution, but such analyses can use finer scale daily data.

In a previous paper, van de Pol et al. [7] provide a broad introduction to climate window analysis for a general scientific audience, with practical details on how the method can be applied using ***climwin***. It proposes a step-wise approach for ***climwin*** implementation that encourages users to identify all potential competing hypotheses, including different potential climate variables (e.g., rainfall, temperature), climate window types (relative or absolute; see Section 1.2), response functions (e.g., linear, quadratic), and aggregate statistics (e.g., mean or maximum temperature). Each of these hypotheses should then be tested and compared using a climate window analysis, with the intention of identifying those hypotheses that are best supported by the data.

This paper is complementary to van de Pol et al. [7], building on the general introduction to ***climwin*** by discussing the technical details of the package, both the design of the package code and the statistical reasoning behind the proposed methods. We discuss a number of topics not covered in the previous paper, including the difference between absolute climate windows (e.g., May to June) and relative climate windows (e.g., two preceding months) (Section 1.2); and the potential use of multi-model inferencing in climate window analysis (Section 1.4). We expand upon the commonly used sliding window analysis, discussed in van de Pol et al. [7], and propose an alternative method for analysing climate, a weighted window analysis (Section 2); we then consider the mechanisms available to account for errors and biases in both methods (Section 3). Finally, we run through a worked example to demonstrate both methods using a real world dataset (Section 4).

In combination, this paper and van de Pol et al. [7] provide a comprehensive overview of ***climwin***; its strengths and weaknesses; and potential future directions for the package. While ***climwin*** has been designed with climate analysis in mind, the package can be applied to any analysis over a continuum (e.g., time or distance) using climatic or non-climatic predictors. For example, climate window methods like those provided in ***climwin*** have been used to analyse plant neighbourhood competition [12]. Therefore, we expect ***climwin*** to have broad applicability both in climate change ecology and more broadly within the scientific community.

## 1 Sliding window analysis

### 1.1 Introduction

#### Model selection metrics

Early sliding window analyses used Pearson’s correlation coefficient to select among different climate window models, where the best window was considered to be the one with the strongest correlation between the climatic predictor and response (e.g., [8–10]). Yet this method only works in simple Gaussian regression models, and there is no possibility to include additional covariates or random effects terms or consider non-linear effects of climate.

Later sliding window studies have used information criteria (IC; [13–15]) as a metric for model selection among competing climate windows (e.g., [5, 11]). An IC-based approach compares all candidate models (i.e. climate windows) and ranks them using a chosen Information Criterion (e.g., Akaike, Bayesian or Deviance Information Criterion; AIC, BIC and DIC respectively). This allows for comparison of any type of multiple regression models, rather than correlation between two variables, and allows users to assess model uncertainty and conduct multi-model inferencing (see Section 1.4). These characteristics make an IC approach more suitable for analysis of climate windows, where it is necessary to compare hundreds or thousands of different models with the aim of determining a best window or group of best windows. An IC approach forms the basis for all climate window comparisons in ***climwin***.

#### Function slidingwin

***climwin*** provides the function `slidingwin` for sliding window analysis. `slidingwin` requires two separate datasets: one containing climate data (ideally at a daily scale) covering the entire period of interest and one containing information on the response variable, as well as any potential covariates. To properly test the relationship between our biological response and climatic predictor, it is necessary for us to take measurements that have different climatic histories. Ideally, this will involve a combination of temporal and spatial replication, where we measure our response variable over multiple years and/or sites. However, combining these two forms of replication assumes that climatic sensitivity is consistent across time and space, which may not always be the case (e.g., [16]).

A key feature of ***climwin*** is the ability for users to define a baseline model into which climate data will be added. This versatility allows for the analysis of data with a variety of error distributions (e.g., Gaussian, binomial, Poisson), the inclusion of multiple covariates, the use of mixed effects modelling, and different types of regression models. Currently ***climwin*** is known to work with base R functions *lm* and *glm* [17], mixed effects model functions from the package ***lme4*** (`lmer`, `glmer`; [18]), and the cox proportional hazard function from package ***survival*** (`coxph`; [19]). Technically, any model that returns a log-likelihood or IC value can be integrated into ***climwin***; however differences in syntax between different modelling packages have hindered our ability to integrate more modelling functions. We aim to provide a greater number of function options for model fitting in future versions.

As highlighted in the introduction, it is possible to vary a broad range of climate window characteristics in `slidingwin` (e.g., temporal resolution of climate data, aggregate statistic, model function). Varying different characteristics of the sliding window analysis allows users to test a variety of climate window hypotheses and help identify potentially novel relationships between climate and the biological response. For example, while we commonly consider mean climate, recent studies have highlighted the potential importance of climatic range [20], rate of climate change [21, 22], and climatic thresholds [23]. However, although it is important to consider a diversity of climate window characteristics in our analyses, changes in many of these characteristics can slightly alter the technical details of the methods used in ***climwin***; therefore, we will focus specifically here on the use of mean climate at a daily resolution.

### 1.2 Relative and absolute climate windows

It is possible that the date of measurement for each record in the response dataset will vary within a sampling group (e.g., year or site). This may be due to constraints on the expression of the response variable (e.g., the date at which offspring size can be measured will depend on birth date) or practical limitations involved in data collection. In cases where the variation in measurement time is small it is reasonable to assume that all records will be influenced by climatic conditions at the same point in time; however, as variation increases this assumption becomes less realistic.

To address this issue, ***climwin*** allows for the use of both absolute and relative climate windows [24, 25]. In an absolute climate window, we assume that all records are influenced by climate at the same absolute point in time, allowing us to define windows using calendar dates (e.g., mean March temperature). Absolute windows require the user to provide a reference date, used as the start point for all fitted climate windows. By contrast, a relative climate window assumes that each record will be impacted by climate at different times depending on the time of measurement. Unlike absolute window analysis, a relative window analysis will test the impact of climate *x* days before the date of measurement.

Absolute climate window analysis is most useful for sampling populations with little temporal variation or data sets where we lack any information on within-group variation in trait expression (e.g., datasets with one aggregate measurement per group; mean body mass of a population). However as temporal variation in the data increases relative windows become more appropriate, particularly when searching for short-lag climate signals. For example, large variation in moult timing of superb fairy wrens (*Malurus cyaneus*) makes the use of an absolute climate window inappropriate as many individuals will already have completed moulting before the start point of the absolute climate window. In this case, a relative climate window (e.g., the 25 days before moulting) is much more useful [25]. It should be noted however, that the output of relative windows can often be more difficult to interpret at the population level as individuals will vary in their climatic sensitivity. Thus the choice of an absolute or relative window involves a trade-off between biological realism and ease of interpretation.

#### Within-group centring

As an absolute window approach assumes no variation in response within a group it can usually only explain between-group variation in the response variable. In comparison, a relative window approach can explain both within- and between-group variation in the response, potentially improving the explanatory power of any fitted climate window model. In certain cases, users may wish to disentangle these within- and between-group climate effects, as they may not necessarily be of equal interest or of the same magnitude. For example, spawning dates of frogs showed a weaker within population response to temperature than that observed across the whole of Britain [26]. ***climwin*** can distinguish both effects by separating climate variables using a technique called within-group centring [27], such that both the within- and between-group climatic sensitivity are estimated for each given time window using the parameter *centre*. Whether one is interested in differentiating between these two types of variation will inform the choice of window type.

### 1.3 How it works

#### Linking climate and biological data

The first step of the `slidingwin` function involves the linking and manipulation of the date information provided in the climate and biological response data frames. As R cannot automatically read date data, ***climwin*** converts this data into an R date format using the function as.Date. Date information must be provided in a standard dd/mm/yyyy format to ensure this process is successful. At this point, we also take into account whether an absolute or relative window is used. Where an absolute window is chosen, the date values of all biological records are changed to the reference day and month provided by the user, with year remaining unchanged.

Using this new date information, `slidingwin` creates a data matrix containing the relevant climate data for each record in the response data frame. For each biological record we extract the climate data needed to fit all potential climate windows (e.g., climate up to 365 days before measurement; Table 1). The amount of climate data stored in this matrix will depend on the minimum and maximum number of days considered in the analysis, determined by the *range* parameter.

**Table 1.**
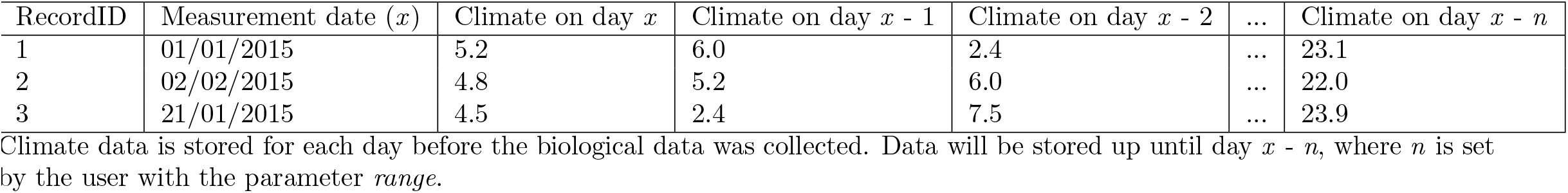
Example of a climate matrix built using *slidingwin*.

#### Model fitting

With a completed matrix we now possess all the necessary information to test different climate windows. `slidingwin` uses nested for-loops to vary the start and end time of climate windows. Where start and end time are acceptable (i.e. start time occurs before end time) `slidingwin` will subset the climate matrix to include only climate data which corresponds to the tested window. We use this data subset to calculate the aggregate statistic (e.g., mean, max, slope), set using the *stat* parameter.

~~~
R> apply(climatematrix, [windowstart:windowend], 1, FUN = stat)
~~~

Where *windowstart* and *windowend* refer to the columns in the climate matrix from which climate data is extracted. The user can decide to test a linear effect of climate, or use more complex model structures (e.g., quadratic, logarithmic, inverse). The function used to test climate is determined by the user with the *func* parameter. Before the for-loops begin, we update the baseline model structure to be consistent with the level of *func*, using a dummy climate variable. Carrying out this structure update before entering the for-loops helps to reduce computational time.

~~~
R> func <- “quad”
R> baseline <- glmer(Response˜ 1 + (1│ID), data = BiolData, family = poisson)
R> BiolData$climate <- rep(1, times = nrow(BiolData))
R> baseline <- update(baseline, .˜. + climate + I(climate^2), data = BiolData)
~~~

Once inside the for-loops, we can replace the dummy climate data with the climate data extracted from the climate matrix. Using the update function we then refit our model.

#### Information criterion

Once we have updated our model to replace the dummy climate data we can extract a sample size corrected measure of AIC (AICc), using the function AICc from the package ***MuMIn*** [28]. However, AICc does not tell us whether a fitted climate window improves upon the baseline model (i.e. a model containing no climate). Therefore, we subtract the model AICc from the AICc value of the baseline model. This creates a metric (ΔAICc) that can be used to both compare individual climate windows to one another and determine how well climate in any given window improves upon the explanatory power of the baseline model. Currently all ***climwin*** functions use AICc as their information criterion; however, there is potential for other criteria to be used in the future.

#### Output

`slidingwin` returns three distinct objects. Firstly, `slidingwin` will return a data frame containing information on the entire model set reflecting all fitted climate windows. This data is sorted by ΔAICc, so that the best model (i.e. smallest ΔAICc value) is listed at the top. With this data frame, the function plotdelta can be used to produce a heat map representing the landscape of ΔAICc values for all fitted climate windows (see Section 4). By examining the ΔAICc landscape the user can determine whether multiple peaks of climatic sensitivity may be present in the data. Additionally, `slidingwin` returns the best model (i.e. the model with the lowest value of ΔAICc) as well as the climate vector used to fit this best model.

### 1.4 Multi-model inferencing

Until this point we have only discussed extracting a single best model from our `slidingwin` analysis; however, we must be aware that there will be uncertainty in the estimation of the best model. An IC approach provides well established methods to deal with this uncertainty, using Akaike model weights (*w_i_*; the probability that model *i* is in fact the best model within the model set; [15]). In practice, we often have little certainty that the model with the lowest ΔAICc is in fact the best model, as a number of top models can have very similar values of *w_i_*. This is particularly likely in climate window analysis as climate data will often be strongly auto-correlated. Our worked examples illustrate that the top models can have very similar values of both ΔAICc and *w_i_* (see Section 4). Is it reasonable, therefore, to extract a single best window from a sliding window analysis?

Ultimately, this will depend on one’s reason for using ***climwin***. Although we often discuss climate as the key point of interest, in some cases users may be more interested in simply accounting for the effect of climate on their response variable, without much concern for the exact nature of the climatic signal. In such a case, it makes sense to extract and use the best climate window as this is, by definition, the climate window that can best explain variation in the response variable.

In other cases, we may be more interested in accurately calculating the timing of a climate signal and/or the relationship between climate and our response. In these scenarios, it makes much less sense to pick a single window as the difference in *wi* between the top windows is likely to be small. As an alternative we can take a group of models that make up a cumulative sum of *wi*. For example, we may group all those models that include the top 95% of *wi*. With such a subset we can be 95% confident that the best model is located within our new model set. This model set is often called a ‘confidence set’ [15]. We can then report values calculated from this subset of top models using multi-model inferencing.

Measuring the percentage of windows included within a confidence set (*C*) can help users determine confidence in a given climate signal. If the models within the set make up a small percentage of the total models tested (*C* is low; e.g., Fig 2a) we can be much more confident that we have observed a real climate signal; however when no climate signal occurs, the confidence set is likely to be much larger (*C* is high; e.g., Fig 2b). ***climwin*** includes the plotting function `plotweights` that visualises different confidence sets for a sliding window analysis and calculates the percentage of models within the 95% confidence set (by default *plotweights* uses the 95% confidence set although users can adjust this cut-off if desired).

**Fig 2.**
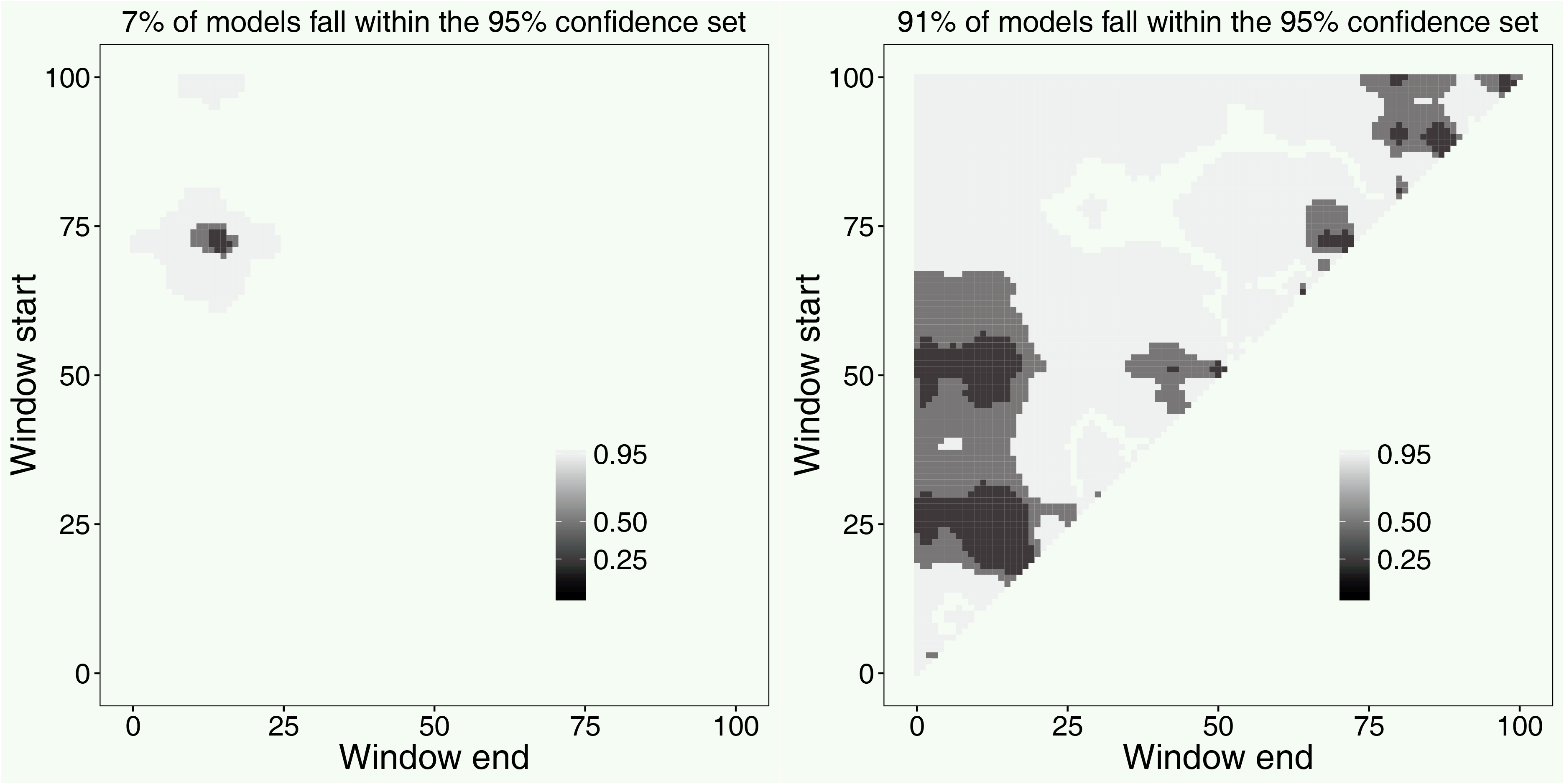
Heat-map of 95%, 50% and 25% confidence sets for slidingwin analysis. Where a strong climate signal occurs, models within the confidence sets make up a small percentage of total models (a; 7%). Where there is no climate signal the confidence set is much larger (b; 91%). A point with window start of 100 and window end of 50 represents a climate window fitted using mean climate 50 - 100 days before measurement date. Figures generated using *plotweights*.

When we are interested in estimating the timing of a climate window, it may be useful to determine a median start and end time for all windows within the confidence set. This can be acheived using the function `medwin`. Additionally, the function `plotwin` can generate box plots illustrating the variation in start and end times. These median values allow users to account for model uncertainty when estimating climate window timing. Similarly, when a user is interested in estimating the relationship between climate and the biological response we can draw information from a subset of potential climate windows using model averaging [15]. A model averaged parameter estimate is simply the sum of parameter estimates weighted by *w_i_*. With such model averaging we can determine the average relationship between climate and our response variable within the confidence set. Users can conduct model averaging using the parameter estimates and model weight values presented in the `slidingwin` output.

Multi-model inferencing is fairly straight forward for datasets with a clear climate signal, where the value of *C* is small, yet this will not always be the case. Large values of *C* may occur when multiple climate signals are present in the data or when the climate signal is weak (i.e. low R^2^), exacerbated by low sample size (Fig 3). Both the median window location and model averaged parameter estimates are less informative in situations where *C* is large as the 95% confidence set may include poor models with spurious parameter estimates [29]. Where multiple peaks are present it can be reasonable for users to adjust the *range* parameter within their `slidingwin` analysis to approach each climate signal separately. However, when a large value of *C* is caused by a weak signal model averaging is not advisable.

**Fig 3.**
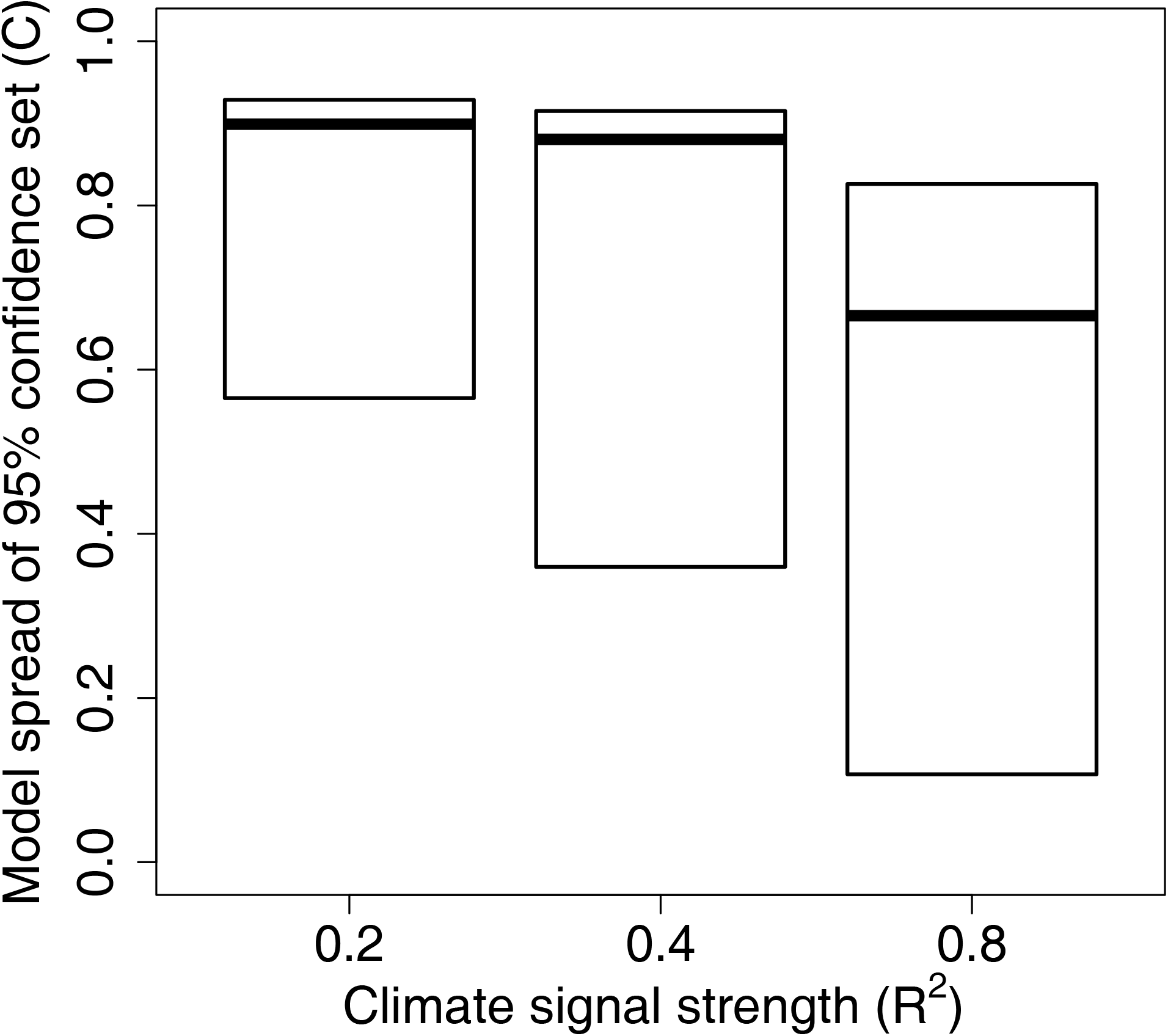
Relationship between the percentage of models in the 95% confidence set and climate signal strength. Percentage of models in the 95% confidence set (*C*) are shown for a very strong (R^2^ = 0.8), strong (R^2^ = 0.4), and moderate climate signal (R^2^ = 0.2). Boxes represent median and inter-quartile range. Data from 2,000 simulated datasets, see Section 3 for methods.

## 2 Weighted window analysis

### 2.1 Introduction

When testing climate windows using mean climate one effectively fits a weight function to the climate data. Using a sliding window approach, we assume that all points between the start and end time of a climate window influence the biological response equally (i.e. a uniform weight distribution with sum of 1). Outside the window, climate is assumed to have no influence on the response (i.e. a uniform distribution with sum of 0; Fig 4a). As we group time into discrete units (i.e. days, weeks, months), assuming a uniform distribution leaves us with a finite number of potential climate windows to test, allowing us to undertake a brute-force approach for climate window analysis, where we systematically test all possible combinations of start and end time sequentially.

**Fig 4.**
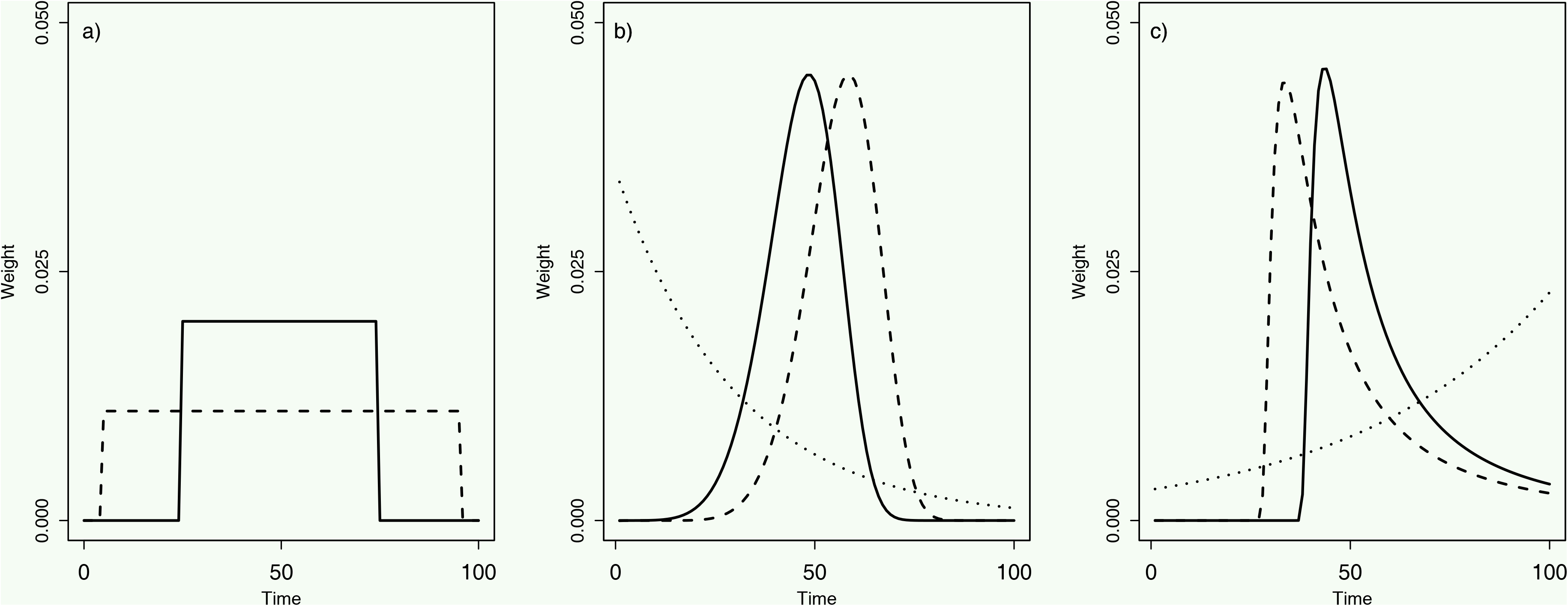
Examples of weight distributions generated with a) uniform, b) Weibull, and c) Generalised Extreme Value probability distribution functions.

Realistically however, the assumption that all points within a time window contribute equally to a climate signal may not be true. The importance of climate will likely change gradually, not abruptly, over time. As an alternative, one can determine a weighted climate mean using a single fitted weight distribution, allowing each climate record to take any weight value between 1 and 0. This allows for more biologically realistic relationships between climate and the biological response. We call this method a ‘weighted window approach’.

***climwin*** includes the function `weightwin`, based on the methods outlined in van de Pol and Cockburn [25], which allows for the calculation of weighted climate means using more complex weight distributions fitted using three parameters: scale, shape and location. The location parameter allows users to adjust where the peak of the distribution sits, similar to a sliding window approach (e.g., Fig 4b solid and dashed lines). Unlike a sliding window analysis however, the scale and shape parameters allow for users to also adjust the width (duration of window) and shape (e.g., exponential decay or bell-shaped) of the distribution respectively. These three parameters are optimised to achieve the lowest possible value of ΔAICc.

As the type of data used is the same, users can apply both the `slidingwin` and `weightwin` function to the same set of data with no changes required. This allows these two approaches to be used in complement to one another and directly compared (section 2.4).

### 2.2 Weight distribution

In principle, any type of probability distribution function can be used to model a weight distribution. So far two probability distribution functions are implemented in `weightwin` that specifically reflect aspects of weight distributions that we think are biologically relevant. The Weibull function is described by the three parameters shape, scale and location and allows for a wide range of weight distributions (Fig 4b). Moreover, for specific values of shape and location the Weibull weight function reduces to an exponential distribution, producing a weight distribution that reflects gradual decay/fading memory effects (Fig 4b; [25]).

The second function is the Generalized Extreme Value (GEV) probability distribution function, which allows for even greater flexibility as it includes functions from the Frechet, Gumbel, and reverse Weibull families (Fig 4c). The GEV function also has a shape, scale and location parameter but, in contrast to the Weibull, includes left-skewed, right-skewed, as well as fairly non-skewed functions, which allows for the comparison of even more refined competing hypotheses. In practice, the GEV function can be harder to fit, as it is more likely to get stuck on local optima during convergence due to the asymptotic nature of the shape parameter around the value zero [25].

Importantly, both the Weibull and GEV probability distribution functions enforce smoothing on the weight distribution. This is of particular importance when analysing climate data, as data is likely to show strong auto-correlation. Furthermore, by imposing smoothing the weight distributions are less likely to be impacted by single extreme climatic events thus reducing the potential for overfitting bias.

### 2.3 How it works

`weightwin` works in a similar way to `slidingwin`. However, rather than varying window start and end time using nested for-loops, `weightwin` varies the values of scale, shape and location to minimise the value of ΔAICc, using the base optimisation function optim in R. By default, we use a quasi-Newton method of optimisation, described by Byrd et al. [30]. This allows for bounding of the shape, scale and location parameters; however, users can employ alternative optimisation methods through the *method* parameter in `weightwin`. Each set of scale, shape and location values is used to generate a weight distribution using either the Weibull or GEV function. This distribution is then used to calculate a weighted climate mean, which is added to the baseline model with the update function. A value of ΔAICc is returned for the optimisation function to assess.

Once the optimisation function has converged, the user will be provided with an output showing the optimised weight distribution and a corresponding best model. Additionally, users will be shown technical details of the optimisation procedure, which can help users to adjust and improve the optimisation process if needed (e.g., alter the initial values with parameter *par* or change the settings of the optimisation routine with parameter *control*).

### 2.4 Comparing approaches

Using a weighted window approach provides a number of benefits over `slidingwin` when assessing the impacts of climate. Firstly, by allowing for an infinite number of potential weight distributions, `weightwin` can provide greater detail on the relationship between climate and the response, such as the occurrence of exponential functions reflecting fading memory effects of past climate. Additionally, by using more diverse weight distributions, `weightwin` will often generate models with better ΔAICc values, which may be especially important when users are most interested in achieving high explanatory power, although one should be aware of potential over-fitting bias (Section 3). Furthermore, by using an optimisation routine `weightwin` often needs to test far fewer models than `slidingwin`, allowing for more rapid analysis.

Despite these benefits, `weightwin` will not always be the most appropriate function for all scenarios. Firstly, the nature of the fitted weight distributions means that `weightwin` can only detect single climate signals, which forces users to detect and compare potential climate signals with separate analyses. While step-wise peak comparison is also required in `slidingwin`, the brute-force approach allows for the detection of multiple climate signals with a single analysis by observing the full ΔAICc landscape. `weightwin` can also be more technically challenging, with users needing to adjust starting values and optimisation settings (e.g., step size, optimisation method) to find the global optimum (i.e. lowest value of ΔAICc). Such technical requirements may limit the accessibility of the `weightwin` function to the general user. Additionally, `weightwin` can only be used for testing mean climate, with no capacity to consider other aggregate statistics. Therefore, whether one chooses to use `weightwin` or `slidingwin` will depend on the aggregate statistic of interest, the level of detail desired, and the user’s technical knowledge.

Ideally, we recommend the use of `slidingwin` and `weightwin` in conjunction to improve our understanding of climate windows. The `slidingwin` approach can be used to explore general trends in the climate data and broadly identify climate signals, including circumstances where multiple climate signals are present. When climate signals are detected using mean climate, the `weightwin` function can then provide greater detail on the specific climate signals observed in the `slidingwin` approach.

### 2.5 Alternative approaches

As discussed above, a limitation of using `weightwin` is the inability to detect and compare multiple climate signals in a single analysis. This issue is a necessary consequence of the assumptions built into the Weibull and GEV functions, forcing us to identify and analyse each climate signal separately. Although `slidingwin` improves upon this issue somewhat by allowing for multiple signal detection, step-wise signal comparison is still required. Yet multiple climate signals may be fairly common and the ability to test and compare these simultaneously would be useful.

With advances in computing and statistics a number of data-driven methods to tackle high-dimensional problems like climate analysis have become common, such as machine learning, least absolute shrinkage and selection operator (LASSO) and functional linear models using splines [12]. These alternative methods offer additional flexibility compared to Weibull and GEV functions, by allowing for the detection of multiple signals with a single analysis (e.g., [12]). Furthermore, they open up the possibility of multi-dimensional climate window analysis, analysing multiple climate variables at the same time, potentially improving upon the uni-dimensional analysis currently employed in ***climwin***.

Splines in particular may provide a suitable alternative for weighted window analysis, as they are ideally suited for modelling a smooth function over a continuum (e.g., time; [12, 31]). In their work, Teller et al. [12] successfully apply a spline function to assess climate signals, demonstrating the ability to detect multiple climate signals within a single weight distribution. Encouragingly, the spline method was able to outperform functions generated by random forest machine learning and LASSO methods, especially at higher climatic resolution that will be common in climate window analyses (e.g., weeks instead of months). The use of splines may reduce the limitations currently encountered by `weightwin`, and incorporating splines is a priority for future ***climwin*** versions.

However it should be noted that the effectiveness of spline functions, in comparison to LASSO and machine learning, was found to vary depending on the characteristics of the data used ([12]; their Fig 6). Users of ***climwin*** will likely analyse a wide variety of data types and seek to answer a broad range of questions. One should be aware that no single method may be ideal for all questions, and it may be more appropriate to consider a range of possible climate window methods and provide a mechanism to compare them. By incorporating a range of alternative methods, such as sliding and weighted window methods, ***climwin*** offers a broad toolbox for analysis of a wide range of questions.

**Fig 6.**
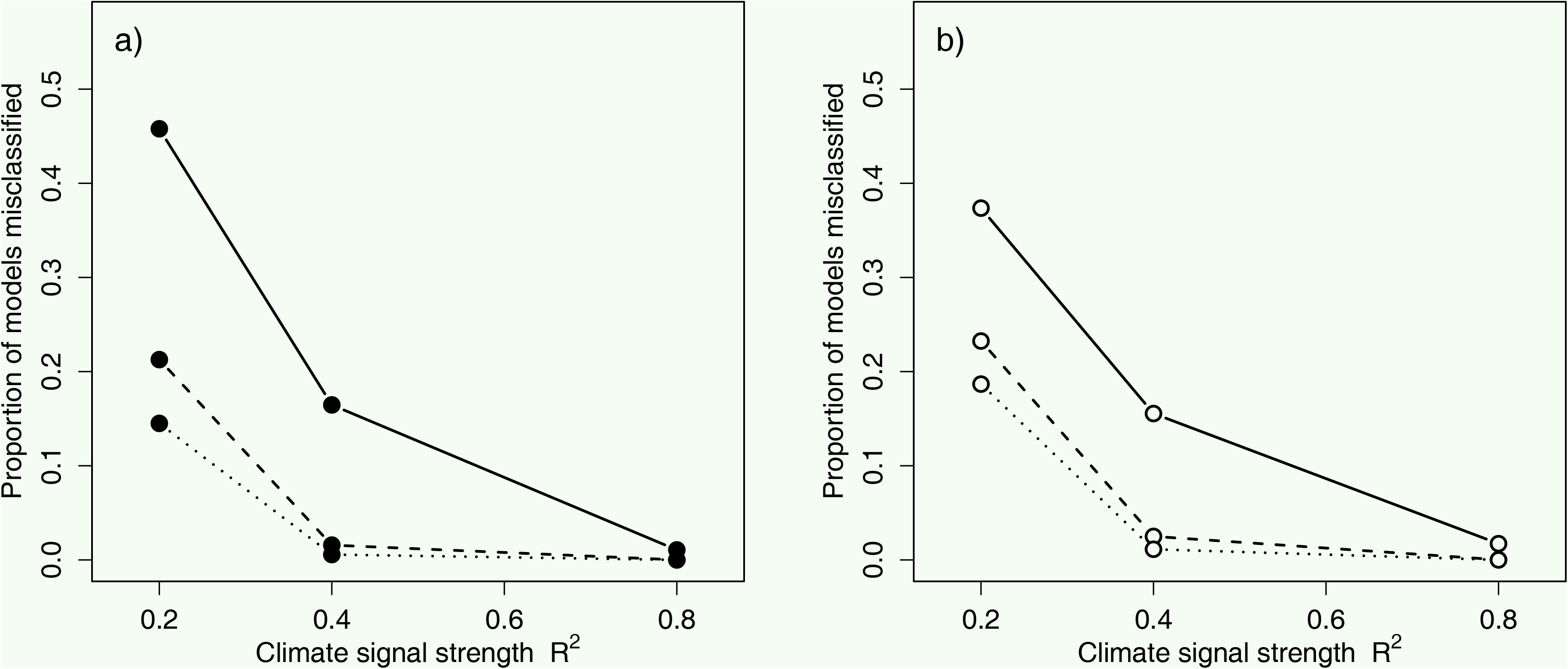
Relationship between climate signal strength (R^2^) and misclassification rate of climate signals. Misclassification rate (false negative) calculated using the metric *P_C_* at sample sizes of 10 (solid line), 30 (dashed line) and 47 (dotted line) with a) no cross-validation and b) 10-fold cross-validation.

## 3 Assessing method performance

Although sliding and weighted window approaches can help us identify climate signals, there has so far been limited systematic testing of the performance of these methods and no way to assess the likelihood that a detected signal is genuine. While Teller et al. [12] employed some method comparison using model correlation (i.e. the correlation of observed parameter estimates with predicted estimates), we still possess little knowledge on potential bias inherent to climate window analyses; the precision of the climate window coefficients and model statistics (e.g., slope, R^2^, window duration); or the rates of type I and type II errors. ***climwin*** includes mechanisms to test and account for many of these potential errors and biases, providing a standard method for testing current and future climate window approaches.

In this section, we will discuss two of these mechanisms, data randomisation and k-fold cross-validation, and quantify their ability to reduce type I and II errors and R^2^ bias respectively. Although we focus here on only two potential biases, users should be aware that biases in other metrics also occur (e.g., slope and window duration bias) and the approaches to account for these biases may differ [7]. Ultimately, the mechanisms one employs to account for potential bias will depend on which metric we most accurately want to predict.

### 3.1 Data randomisation

To estimate the probability that a given result represents a false positive (type I error) we can calculate the expected distribution of ΔAICc values in a data set where no relationship exists between climate and our response variable. ***climwin*** provides the function randwin, which randomises a given dataset (i.e. removes any climate signal) and conducts a sliding window analysis to extract a value of ΔAICc. *randwin* reorders the date variable in the original response data frame, allowing us to maintain any relationship between the response variable and other covariates and maintaining auto-correlation within the climate data while still removing any relationship between climate and the response. Following this randomisation procedure, *randwin* will run a climate window analysis on this new set of data from which we extract the ΔAICc of the best model.

The randomisation process is repeated a number of times, defined by the user with the parameter *repeats*. We recommend a large number of randomisations (e.g., 1,000) to best estimate the distribution of ΔAICc values that could be obtained from a climate window analysis on a dataset with no climate signal (ΔAICcrand). We can then determine the percentile of ΔAICc_rand_ that exceeds the value of ΔAICc observed in our analysis, allowing us to calculate the likelihood that a given ΔAICc value might occur by chance (termed *P*_Δ*AICc*_). *P*_Δ*AICc*_can be obtained using the function pvalue.

Although conducting a large number of randomisations is the best method to guard against false positives, running this many randomisation can be impractical. Many analyses will use large datasets and/or complex models that can take multiple hours to run. Running time will also be impacted by the range over which the analysis covers, with the number of models run during a sliding window analysis increasing approximately quadratically with analysis range (Eq. 1)

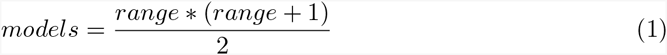

For a sliding window analysis covering a year (`range = c(365, 0)`) ***climwin*** will fit over 67,000 models.

Consequently, carrying out 1,000 or even 100 randomisations may simply take too long for many users. Yet it is still important that we are able to protect against the possibility of false positives. As an alternative, ***climwin*** includes a metric that can be used to estimate the probability of false positives with a limited number of randomisations (e.g., 5 - 10).

To empirically derive an alternative metric, we analysed a range of simulated datasets where the occurrence of a real signal was known. We generated groups of 2,000 datasets, each with a range of sample sizes (10, 20, 30, 40, or 47 datapoints) and levels of climate signal strength (climate signals with an R^2^ that was very high [0.80], high [0.40], moderate [0.20], or where no signal was present). Our simulated datasets were intentionally small, which allowed us to derive a potential metric that is able to function well in challenging situations. Many climate analyses will use datasets with many more data points by employing temporal and spatial replication. The performance of the metric will often be much better in these circumstances.

We assigned each dataset a binary value (*SignalTrue*) depending on whether it contained a real signal (1) or no signal (0). For every dataset, we then ran a full `slidingwin` analysis and extracted metrics for the best model, here after termed the observed result (R^2^, sample size, ΔAICc, and the percentage of models within the 95% confidence set [*C*]). In addition, we ran each dataset either with k-fold cross-validation (with *k = 10* folds; see Section 3.2) or without. In total, we tested 80,000 different datasets. For each of these datasets we then used *randwin*, with *repeats = 5*, to determine the median value of ΔAICc and *C* from randomised data. From this we calculated two new metrics:

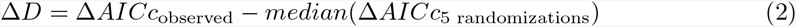

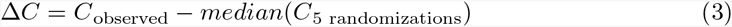

We divided our simulation results in half to generate a training and test dataset that we could use to calculate our new metric. We expected that the effectiveness of Δ*D* and Δ*C* would vary with both sample size and the use of cross-validation. We therefore divided our training dataset again to separate those datasets that used cross-validation and those that didn’t. For each of these two training datasets we then fitted two potential models:

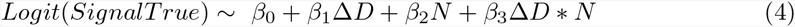

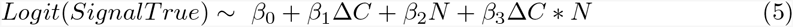

where *N* is the sample size of the dataset used to calculated the values of Δ*C* and Δ*D*.

Both with and without cross-validation, Eq. 5 was clearly the best supported (ΔAICc <-2,500), suggesting that Δ*C* is the best metric to determine the likelihood of a real signal. Therefore, we determine the likelihood that a given value of ΔAICc has occurred by chance with our new metric (*P_C_*) to be:

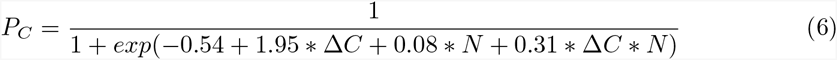

for datasets analysed with the use of 10-fold cross-validation, and

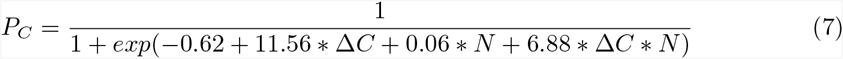

for datasets analysed with the use of 10-fold cross-validation.

Finally, we used our test dataset to determine the rate of misclassification for our new metric, *P*_*C*_. Specifically, we calculated the rate of false negatives in datasets where we knew a signal was present and the rate of false positives in those datasets where no signal existed.

*P*_*C*_ was able to provide a good estimate of the reliability of a signal, with average rates of misclassification generally low (Fig 5; mean false negative rate = 0.10, mean false positive rate = 0.17). The effectiveness of *PC* was strongly influenced by both sample size (Fig 5) and climate signal strength (Fig 6), with misclassification rates dropping well below the overall average when sample size and signal strength increased (e.g., false negative rate = 0.02 when *N* = 30, R^2^ = 0.4; Fig 6). Sample size also had a strong influence on false positive rates which decreased with increasing sample size (Fig 5b). These results are not necessarily surprising as misclassification is common when dealing with weak effects and small sample sizes, but it highlights the importance of using large sample sizes when conducting these types of exploratory analyses and the need for caution when interpreting results from small datasets.

**Fig 5.**
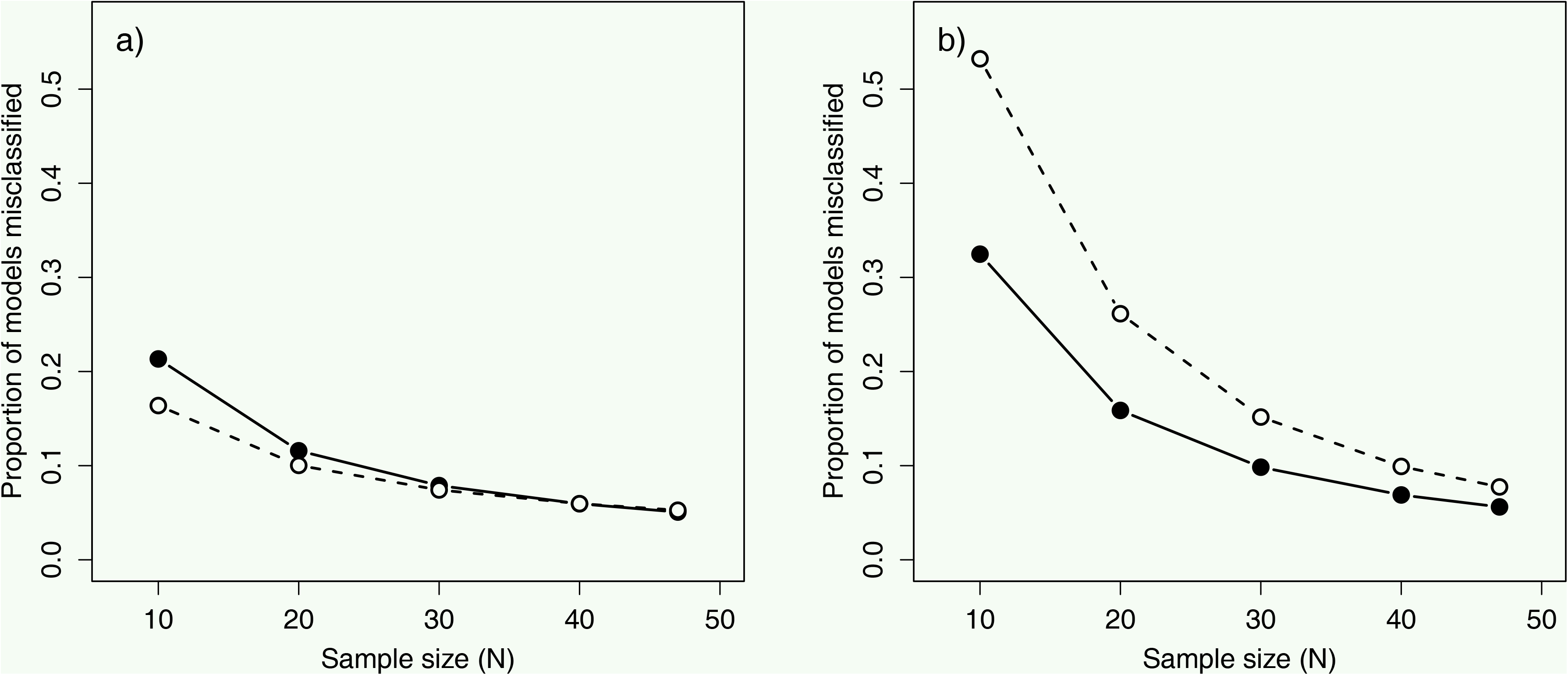
Relationship between sample size (*N*) and misclassification rate of climate signals. Misclassification rate calculated using the metric *P*_*C*_ both with 10-fold cross-validation (dashed line) and without cross-validation (solid line). Metric tested on datasets where a) a climate signal is present and b) a climate signal is missing. Note that misclassification in a) denotes false negatives while in b) it denotes false positives.

For this exercise, we considered a signal to be identified when *P*_*C*_ < 0.5 (i.e. when *P*_*C*_ calculated that there was a better than even chance that a given signal was real). The point that one chooses to distinguish between real and false signals will ultimately involve a trade-off between false positive and negative rates. A lower more conservative cut-off would reduce the chance of false positives but simultaneously increase false negative rates. As an alternative to cut-off values, we encourage the reporting of the full values of *P*_*C*_ and *P* Δ*AICc* as a means of documenting the confidence in a given result, rather than trying to classify signals as either real or not.

### 3.2 k-fold cross-validation

While *P*_*C*_ and *P*_ΔAICc_ can help test the rates of false positives and negatives, they give us no indication of the reliability of the parameter estimates and model statistics derived from our best model (e.g., R^2^, slope, window duration). k-fold cross-validation, provided in `slidingwin`, can be a key tool to help account for any potential biases in these estimates that might arise from overfitting [32]. k-fold cross-validation involves the division of a dataset into *k* training datasets (of length 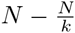) and *k* test datasets (of length 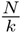, with *k ≤ N*), where *N* represents sample size. Once these training and test datasets are partitioned, `slidingwin` fits each climate model to one of the training datasets and its predictive accuracy is then tested on the corresponding test dataset. To measure predictive accuracy, mean square error (MSE) of the training fit to the test data is used to calculate the AIC_c_:

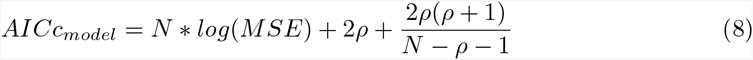

(where *ρ* is the number of estimated model parameters) and subsequently compared to the AICc of the baseline model, also determined using the training dataset, to obtain ΔAICc_model_. This procedure is repeated *k* times (once for each test dataset), after which the ΔAICc_model_ is averaged across all folds to obtain the cross-validated ΔAICc _model_. The total number of folds used, is set by the user with the parameter *k* in the `slidingwin` function.

Cross-validation is used in `slidingwin` to improve the ΔAICc predictions of each climate window, the out-of-sample ΔAICc, which is then used to improve the model selection process. Each climate window is ultimately fitted to the full dataset, so all other parameter estimates and model statistics (e.g., R^2^) have not been cross-validated. However, our more conservative model-selection process is able to greatly reduce the bias in the estimation of climate signal R^2^, reducing the inherent optimistic bias observed in climate window analyses conducted without cross-validation (Fig 7).

**Fig 7.**
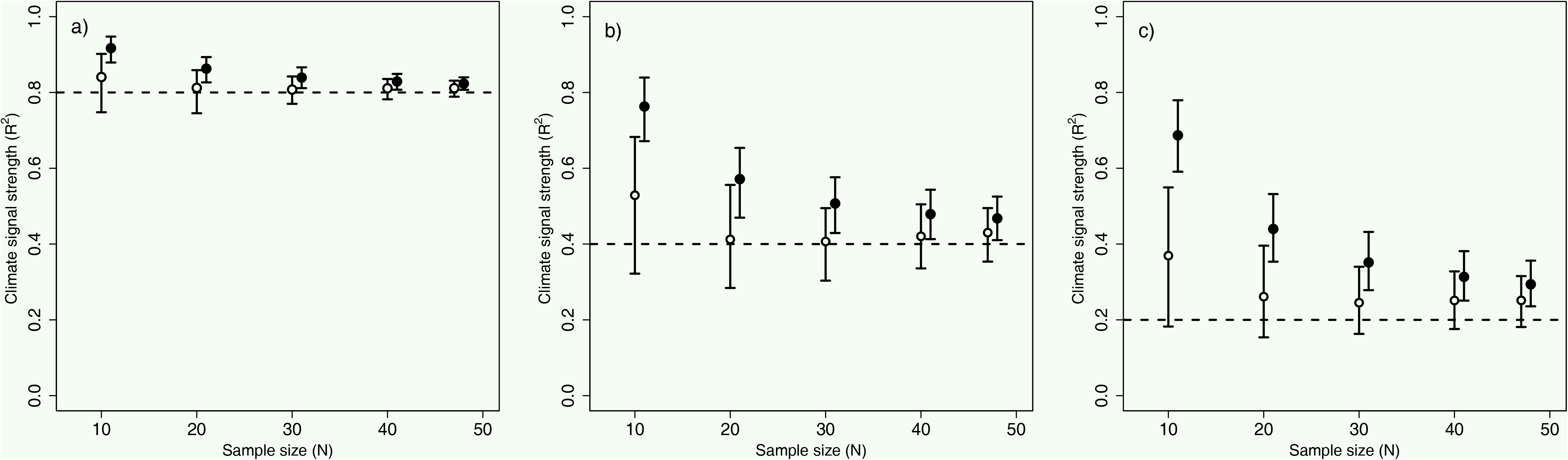
Performance of slidingwin in estimating the true R^2^value of a climate signal. Performance determined at varying sample sizes with very high R^2^ (0.80; top left), high R^2^ (0.40; top right), and moderate R^2^(0.20; bottom) both without cross-validation (black) and with 10-fold cross-validation (white). Points represent median R^2^estimates from 2,000 simulated datasets. Error bars represent inter-quartile range. The horizontal dashed line shows the true value of R^2^ used to generate the simulated datasets.

To determine the optimum value of *k* for R^2^ estimation, we generated groups of 1,200 datasets each with a known climate signal (R^2^ = 0.22) and varying sample sizes (10, 20, 30, 40, or 47 datapoints). For each sample size group, `slidingwin` analysis was conducted varying the value of *k* (0, 2, 4, 6, 8, and 10-folds), so that 200 datasets were tested for each level of sample size and k-folds. Because *k* cannot exceed *N*, *k* = 10 was used as the largest number of folds. We found that increasing the number of folds consistently improved estimation of R^2^ across all sample sizes, with *k* = 10 providing the best estimate of R^2^ (Fig 8).

**Fig 8.**
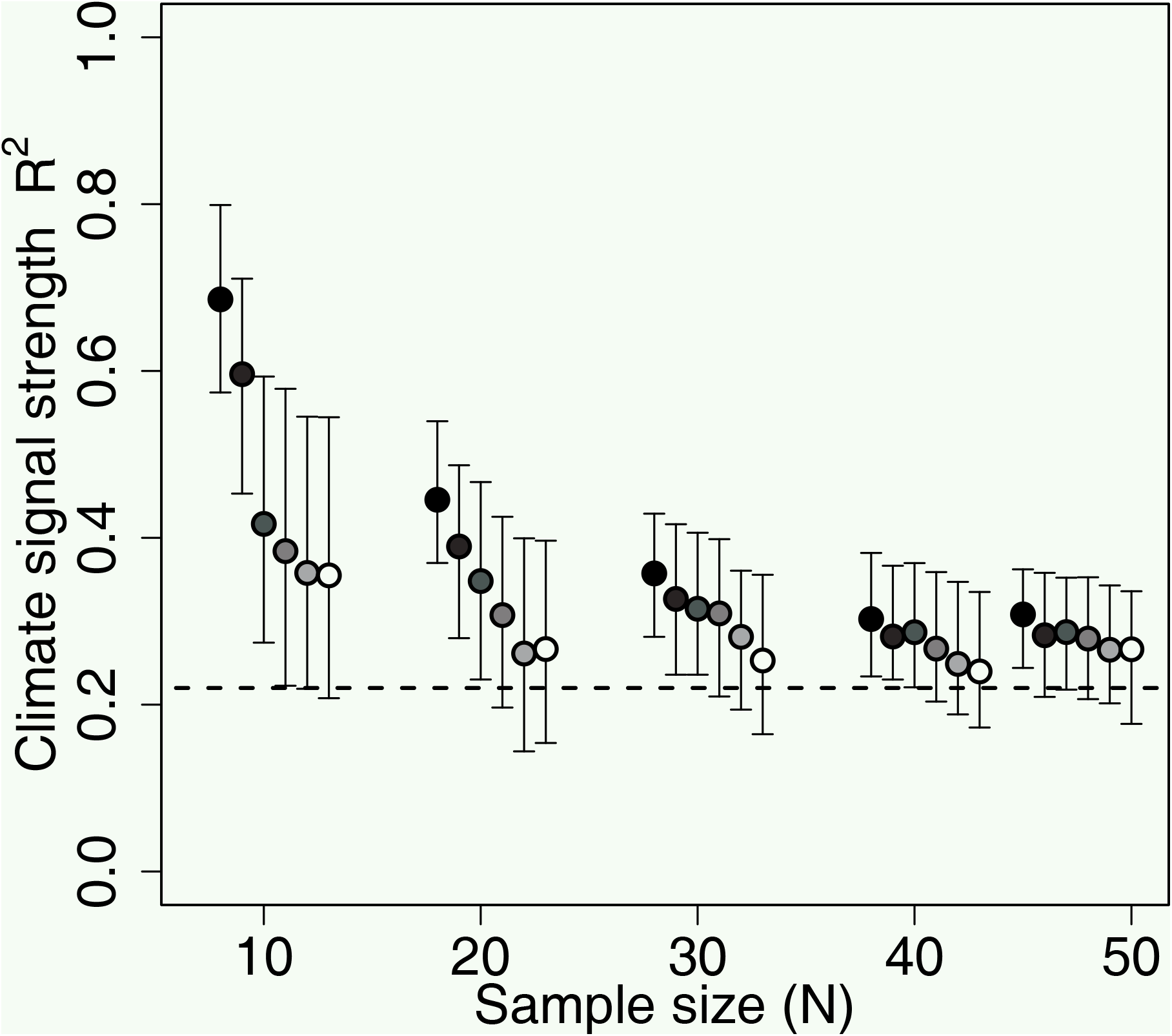
Effect of cross-validation folds (k) on the median R^2^estimation of k-fold cross-validated *slidingwin* analysis. Data generated using 200 simulated datasets. The horizontal dashed line shows the true value of R^2^ used to generate the simulated datasets (R^2^ = 0.22). R^2^ was estimated using 0, 2, 4, 6, 8, or 10-folds (black to white respectively). Sample sizes of 10, 20, 30, 40, and 47 were used. Error bars represent inter-quartile range.

Although cross-validation greatly improves R^2^ estimation, users should be aware that R^2^ bias is not completely removed by cross-validation and the goodness-of-fit of the best model from `slidingwin` may still be overly optimistic. Additionally, like data randomisation, k-fold cross-validation can substantially increase the computational time of `slidingwin`, and users will need to consider a trade-off between reducing R^2^ bias and analysis time.

While data randomisation and k-fold cross-validation improve our detection of climate signals and our estimates of climate signal R^2^, neither of these methods can be reliably used to simultaneously combat all potential biases in climate window analysis. For example, although cross-validation can effectively reduce bias in R^2^ it will also increase false positive rates, particularly at low sample sizes (Fig 5b). Ultimately, therefore, the methods chosen to reduce bias in climate window analysis will differ depending on the particular parameters of interest.

## 4 Worked examples

This section provides examples applying the ***climwin*** package to real data. We use the `Chaff` and `ChaffClim` datasets, included with the package, to run both a sliding window and weighted window analysis. As part of this analysis, we demonstrate the use of multi-model inferencing to determine the median start and end time of a climate signal and conduct model averaging on parameter estimates. In addition, we conduct k-fold cross validation and data randomisation to determine *P*_*C*_ and *P*_Δ*_AICc_*_.

### 4.1 Analysis with slidingwin

Our analysis of the `Chaff` dataset focuses on the impact of mean temperature on the annual average laying date of the common chaffinch (`Fringilla coelebs`) over a 47 year period (1966-2012; with data provided by the British Trust for Ornithology). We first carry out a sliding window analysis on our data using `slidingwin`.

#### Function syntax

To begin, we set the structure of our baseline model using the base *lm* function.

~~~
R> baseline = lm(Laydate˜ 1, data = Chaff)
~~~

Although we use a simple baseline model for illustration, it is possible to include covariates and random effects terms into the baseline model, as well as using different model functions (e.g., *lmer*, *coxph*). We next specify the climatic variable of interest using the parameter *xvar* (xvar = list(Temp = Chaff$Temp)), and include both the climate and biological date data with the parameters m*cdate* and `bdate` (`cdate = ChaffClim$Date, bate = Chaff$Date`). As our Chaff dataset contains no within-year variation, we conduct our analysis using absolute climate windows (type = ^“^absolute^”^) with a reference day of April 24th (`refday = c(24, 4)`), equivalent to the earliest biological record in our data.

As we have no `a priori` knowledge on when a climate signal might occur, we test all possible climate windows over the period of a year (`range = c(365, 0)`), considering the linear effect (`func = “lin”`) of mean temperature (`stat = “mean”`). With all these elements, our final function is shown below:

~~~
R> SLIDING <- slidingwin(baseline = lm(Laydate˜ 1, data = Chaff),
                  Xvar = list(Temp = ChaffClim$Temp),
                  cdate = ChaffClim$Date, bdate = Chaff$Date,
                  type = “absolute”, refday = c(24, 4),
                  range = c(365, 0), func = “lin”, stat = “mean”)
~~~

By default, `slidingwin` will assume daily climate data is used to test climate windows. However, in cases where the resolution of climate data is coarser, users can alter the parameter cinterval to use either weeks or months.

## Results

The object `SLIDING` is a list item with two separate elements. We can firstly examine a summary of our results using the `combos` item, a truncated version of which can be see in Table 2.

**Table 2.**
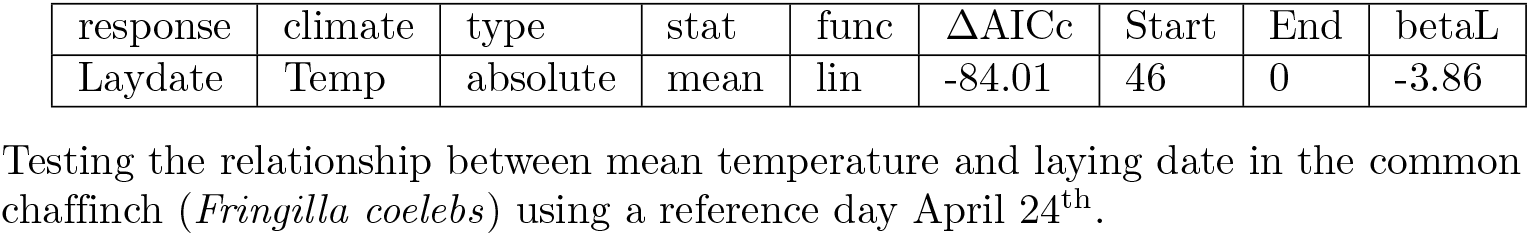
Output of *combos* item from an absolute sliding window analysis.

| response | climate | type | stat | func | $\Delta AICc$ | Start | End | betaL |
| --- | --- | --- | --- | --- | --- | --- | --- | --- |
| Laydate | Temp | absolute | mean | lin | -84.01 | 46 | 0 | -3.86 |

Testing the relationship between mean temperature and laying date in the common chaffinch (*Fringilla coelebs*) using a reference day April 24<sup>th</sup>.

R> SLIDING$combos

The `combos` item provides a summary of our sliding window analysis and a brief overview of the best fitted climate window, showing us the ΔAICc, start and end time, and slope of the best window. It should be noted that ***climwin*** allows for multiple hypotheses to be tested in a single function (e.g., effect of mean and maximum temperature), in which case the combos item will provide a summary of all tested hypotheses. For this example, we can see that the best climate window detected in our analysis falls 46-0 days before our reference date (April 24^th^), equivalent to mean temperature between March 9^th^ and April 24^th^.

We can look at the results further in the full model selection dataset, a truncated version of which can be seen in Table 3.

**Table 3.**
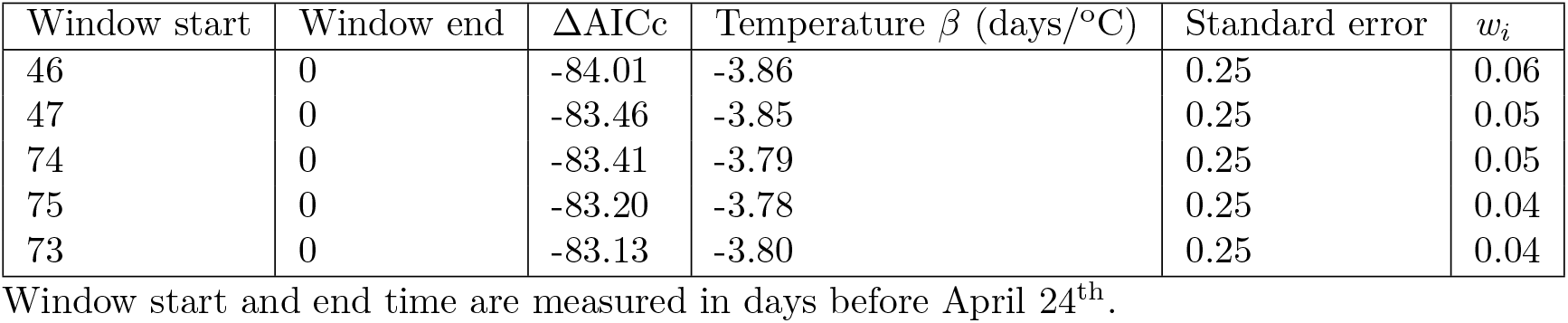
Top five climate windows detected using *slidingwin* with an absolute window approach.

~~~
R> head(SLIDING[[1]]$Dataset)
~~~

In Table 3 we can see that there are a number of climate windows that exhibit similar model weights (*w_i_*) to our best window. To understand how these other windows influence our result we can determine the median window size of the 95% confidence set with our function `medwin` and calculate model averaged parameter estimates for the same confidence set.

~~~
R> medwin(SLIDING[[1]]$Dataset)
R> dataset <- SLIDING[[1]]$Dataset
R> ConfidenceSet <- dataset[which(cumsum(dataset$ModWeight) < = 0.95), ]
R> sum(ConfidenceSet$ModelBeta*ConfidenceSet$ModWeight)
~~~

Median window size from the 95% confidence set is slightly wider than our best window (73 - 1; February 11^th^- April 23^rd^), although the median and best window still contained over 60% of the same days. The best window shows a strongly negative relationship between temperature and laying date (*β* = -3.86 days/^o^ C, 95% CI = -4.35 - -3.37; Table 3), very similar to the model averaged relationship (*β* = -3.60 days/o C). Multi-model inferencing tells us that the average laying date of *F. coelebs* advances by 3.6 days for every 1o C increase in mean temperature between February 11^th^ and April 23^rd^.

Although these results point to the presence of a strong climate signal in *F. coelebs* laying date, we cannot be sure that this result has not occurred due to chance. To test this possibility, we next run the randomisation procedure using the function randwin, with repeats = 5.

~~~
R> SLIDING.RAND <- randwin(repeats = 5,
                                         baseline = lm(Laydate˜ 1, data = Chaff),
                                         xvar = list(Temp = ChaffClim$Temp),
                                         cdate = ChaffClim$Date, bdate = Chaff$Date,
                                         type = “absolute”, refday = c(24, 4),
                                         range = c(365, 0), func = “lin”, stat = “mean”>)
~~~

The output of the randwin function can then be used to run the function pvalue to return a value of *P*_*C*_.

~~~
R> pvalue(dataset = SLIDING[[1]]$Dataset,
         datasetrand = SLIDING.RAND[[1]], metric = “C”>, sample.siz         e = 47)
~~~

From this function, we can conclude that the likelihood of observing such a climate signal by chance is very small (*P*_*C*_ = 5.89e^-16^).

Although this provides us with information on the best model, it does not tell us whether multiple peaks may be present. Our final step should therefore be to examine the ΔAICc and model weight landscape (Fig 9). In this case, there is only a single clear ΔAICc peak (red; Fig 9a), which is mirrored in the small size of the confidence set (*C*) (Fig 9b). We can therefore discount the possibility of multiple peaks.

**Fig 9.**
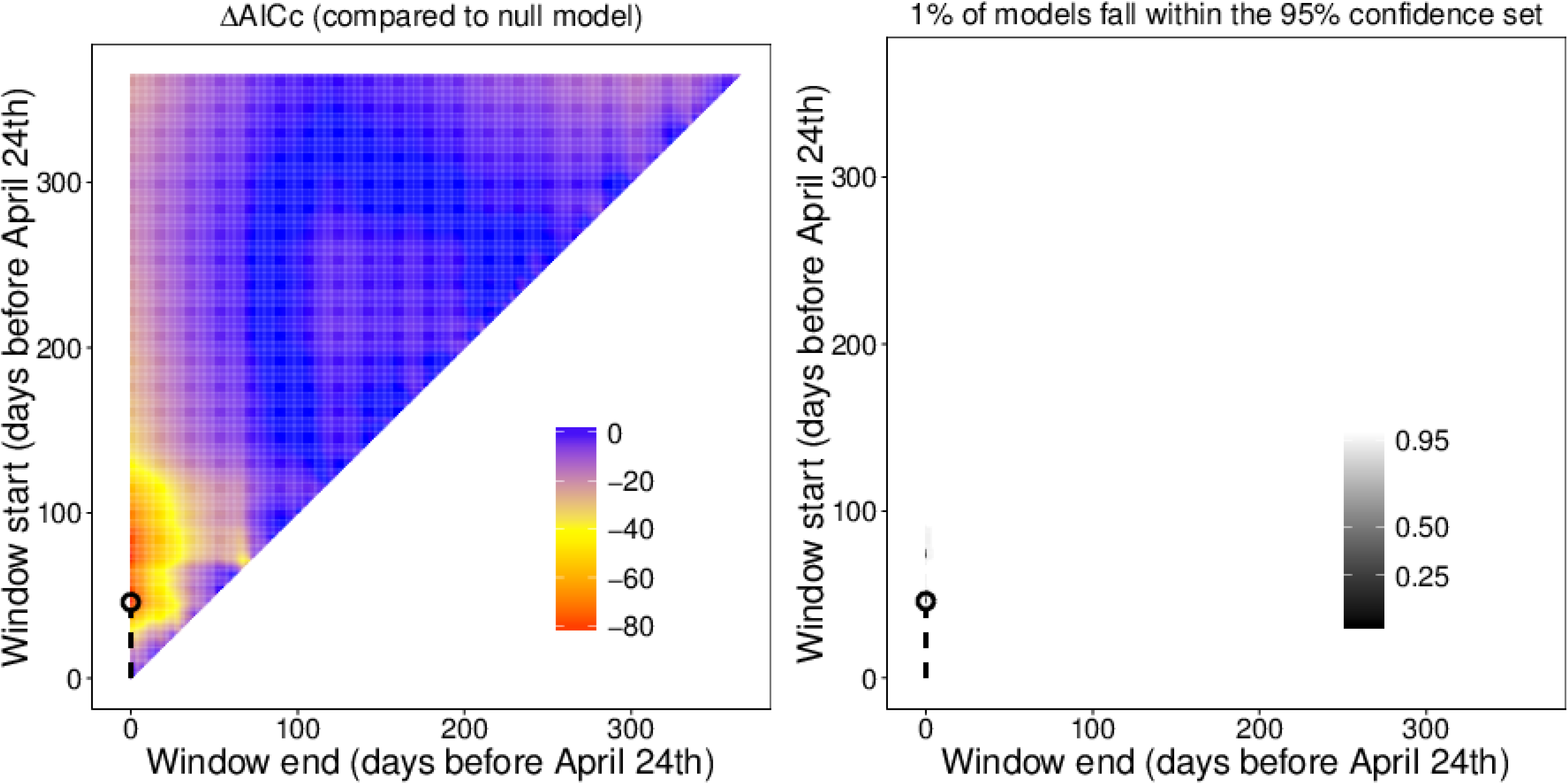
Output of absolute sliding window analysis. Analysis testing the relationship between mean temperature and laying date in the common chaffinch (`Fringilla coelebs`) using a reference day April 24^th^. (*Left*) Heat map of ΔAICc (AICc of null model - AICc of climate model) for all fitted climate windows. (*Right*) 95%, 50% and 25% confidence sets for all fitted climate windows. The best fitted climate window (lowest value of ΔAICc) is circled. Plots generated using plotdelta and plotweights functions.

### Using k-fold cross-validation

Above, we have focused on estimating the window duration and slope using multi-model inferencing. However, in other circumstances we may be more interested in determining the strength of the detected climate signal (R^2^). As R^2^ estimations using `slidingwin` can be biased at low sample size and/or effect size, k-fold cross-validation should be employed to improve the accuracy of our R^2^ estimate. To conduct our `slidingwin` analysis with k-fold cross-validation we incorporate the parameter k into the `slidingwin` function (k = 10).

~~~
R> SLIDINGK <- slidingwin(baseline = lm(Laydate˜ 1, data = Chaff),
                    xvar = list(Temp = ChaffClim$Temp),
                    cdate = ChaffClim$Date, bdate = Chaff$Date,
                    type = “absolute”, refday = c(24, 4),              range = c(365, 0), func = “lin”, stat = “mean”, k = 10)
~~~

Looking at the combos object, we can see that the best model selected using cross-validation has a very similar window duration and slope to that calculated using multi-model inferencing in our first sliding window analysis (Window duration: 75 - 0, February 9^th^ - April 24^th^; window slope: -3.78 days/^o^ C, 95% CI = -4.27 - -3.30; Table 4).

**Table 4.** Output of *combos* item from an absolute sliding window analysis.

| response | climate | type | stat | func | ΔAICc | start | end | betaL |
| --- | --- | --- | --- | --- | --- | --- | --- | --- |
| Laydate | Temp | absolute | mean | lin | -11.07 | 75 | 0 | -3.78 |

Testing the relationship between mean temperature and laying date in the common chaffinch (*Fringilla coelebs*) using a reference day April 24<sup>th</sup> and 10-fold cross-validation.

`R> SLIDINGK`$combos

Although window duration and slope are similar to our previous analysis, the value of ΔAICc is much less negative, due to the conservative nature of ΔAICc calculation when using cross-validation (i.e. ΔAICc is calculated on a smaller test dataset). This more conservative ΔAICc estimation will also lead to much larger values of *C* (Fig 10), which will often remove the possibility for users to conduct multi-model inferencing. However, even though the model weight landscape shows less compelling evidence of a climate signal, by running `randwin` with cross-validation and calculating *PC*, we find that the likelihood of getting such a value of *C* by chance when using 10-fold cross-validation is still very small (*P*_*C*_ = 1.10e^-11^).

**Fig 10.**
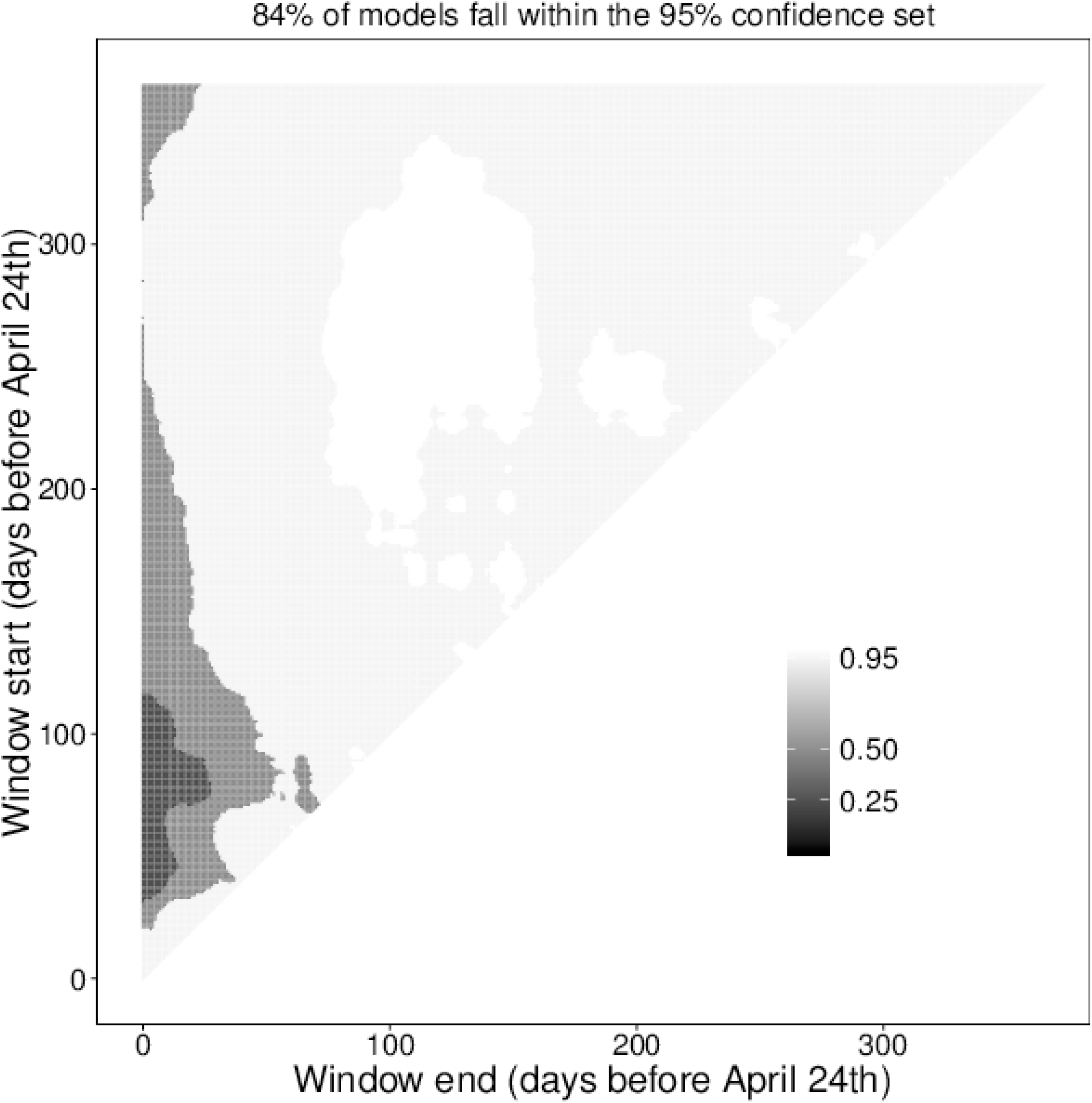
Heat-map of 95%, 50% and 25% confidence sets for an absolute sliding window analysis. Analysis testing the relationship between mean temperature and laying date in the common chaffinch (`Fringilla coelebs`) using a reference day April 24th and 10-fold cross-validation. Shading levels represent 95%, 50% and 25% confidence sets for all fitted climate windows. Plots generated using the plotweights functions.

Once we are confident in our climate signal result we can then examine the summary of the best model to gain an estimate of strength for the climate signal.

~~~
R> summary(SLIDINGK[[1]]$BestModel)
~~~

In this case, the strength of the climate signal detected in *F. coelebs* laying date is particularly strong (R^2^ = 0.83).

### 4.2 Analysis with weightwin

Using `slidingwin` we have been able to identify a negative relationship between mean temperature and *F. coelebs* laying date. Yet we have so far assumed a uniform weight distribution when calculating mean temperature. To gain more insight into the detected climate signal, we can next run a weighted window analysis using `weightwin`.

Firstly, we want to determine the best starting distribution to use for the `weightwin` optimisation procedure, using the included `explore` function. We can experiment with the shape, scale and location parameters for a Weibull distribution to determine a reasonable starting weight distribution for our optimisation procedure (Fig 11).

**Fig 11.**
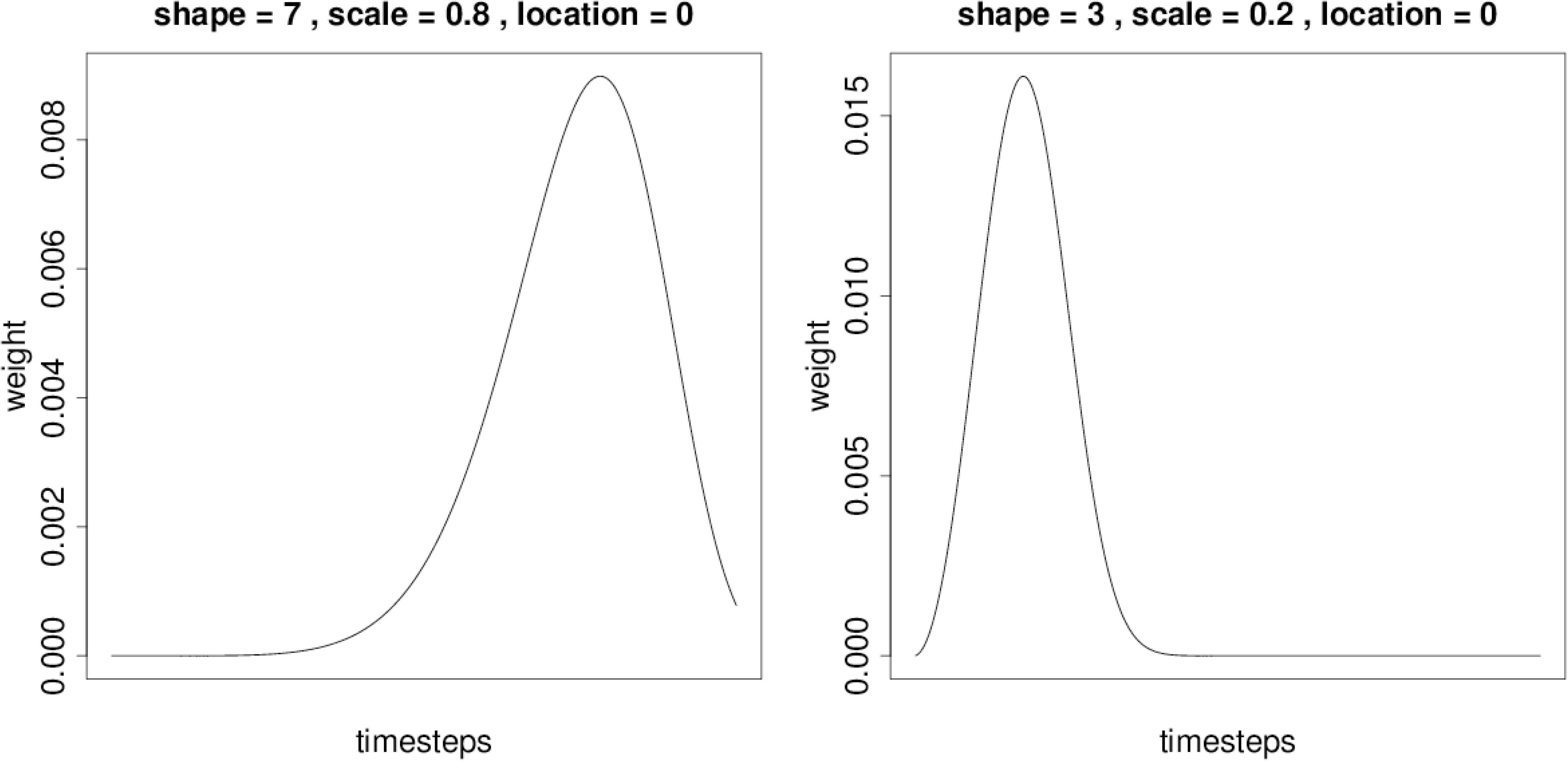
Weight distribution calculated using a Weibull probability distribution function. Distribution shows the relative importance of climate over time (days). (*Left*) Values of shape, scale and location used as starting parameters for weighted window analysis. (*Right*) Output from `weightwin` analysis showing the relative influence of temperature on the average annual laying date of the common chaffinch (`Fringilla coelebs`). Weight distribution shows that temperature has the strongest influence on laying date immediately before the reference date (April 24^th^) but slowly decays as we move further into the past. Plots created using the function explore.

~~~
R> explore(shape = 3, scale = 0.2, loc = 0, weightfunc = “W”)
~~~

Most of the parameter values will be the same between `weightwin` and `slidingwin`, but we must provide additional information on the type of probability distribution function being used (in this case Weibull, `weightfunc` = “W”) and the starting values of our three optimisation parameters, taken from the explore function (par = c(3, 0.2, 0)). Additionally, both the parameters *k* and *stat* are not used in `weightwin`.

~~~
R> WEIGHT <- weightwin(baseline = lm(Laydate˜ 1, data = Chaff),
                   xvar = list(Temp = ChaffClim$Temp),
                   cdate = ChaffClim$Date, bdate = Chaff$Date,
                   type = “absolute”, refday = c(24, 4),
                   range = c(365, 0), func = “lin”,
                   weightfunc = “W”, par = c(3, 0.2, 0))
~~~

In contrast to the uniform distribution assumed by `slidingwin`, our analysis with `weightwin` returned a rapidly decaying weight distribution, with temperature having the largest impact on laying date close to April 24^th^ and rapidly declining further into the past (Fig 11). Furthermore, by examining the WeightedOutput item generated by `weightwin`, we can see that the explanatory power of this weight distribution (ΔAICc) is much greater than that generated with the uniform distribution assumption in `slidingwin` (-84.01 v. -100.42; Table 5).

**Table 5.** Output of an optimised weight distribution (Weibull function) testing the relative influence of temperature on the laying date of the common chaffinch (*Fringilla coelebs*).

| $\Delta\text{AICc}$ | shape | scale | location | model $\beta$ | standard error |
| --- | --- | --- | --- | --- | --- |
| -100.42 | 2.99 | 0.32 | -0.26 | -4.28 | 0.23 |

~~~
R> WEIGHT$WeightedOutput
~~~

Once again, however, we cannot be sure that such a result could not occur by chance and so we can compare our result to those from a randomised dataset using `randwin`.

In this case, however, the smaller computational time required to run `weightwin` allows us to increase repeats to 1,000. Note, however, that we must specify we are running a weighted window analysis with the argument `window` = “`Weighted`”.

~~~
R> WEIGHT.RAND <- randwin(repeats = 1000, window = “weighted”,
                    baseline = lm(Laydate˜ 1, data = Chaff),
                    xvar = list(Temp = ChaffClim$Temp),
                    cdate = ChaffClim$Date, bdate = Chaff$Date,
                    type = “absolute”, refday = c(24, 4),
                    range = c(365, 0), func = “lin”,
                    weightfunc = “W”, par = c(3, 0.2, 0))
~~~

With 1,000 randomisations, we are able to use the more reliable *P*Δ*AICc* to estimate the probability that we would observe such a largely negative value of ΔAICc by chance.

~~~
R> pvalue(dataset = WEIGHT$WeightedOutput,
         datasetrand = WEIGHT.RAND[[1]], metric = “AIC”)
~~~

Once again, we find that the probability of observing such a weight distribution by chance is very small (*P*_Δ*AICc*_ < 0.001). Therefore, our analysis using ***climwin*** provides good evidence that laying date in *F. coelebs*is strongly impacted by temperature over late winter and early spring (February - April) with a decaying relationship over time.

### 4.3 Replication

The worked examples above can be replicated using functions and data included with ***climwin***. The full release version of ***climwin*** (version 1.0.0) is available from the Comprehensive R Archive Network at http://CRAN.R-project.org/package=climwin. The current pre-release version of the package can be accessed on GitHub https://github.com/LiamDBailey/climwin. The worked examples above use the Chaff and ChaffClim datasets included with the full release version of the package. All code was written by Liam D. Bailey and Martijn van de Pol and can be used freely according to the General Public License (GPL), version 2.

## 5 Conclusion

The way in which previous research has tested and compared the effects of climate has tended to require arbitrary *a priori* selection of a limited number of climate windows, curtailing our ability to make meaningful conclusions. Climate window analyses, such as sliding and weighted window analyses, improve on these methods by reducing the need for *a priori* assumptions. Yet until now, we have lacked a standardised and accessible way in which to carry out such analyses, nor any way to assess method performance. We introduced the R package ***climwin***, which provides an easy and versatile toolbox for analysing the impacts of climate using a number of potential methods and includes metrics to assess the performance of these methods. This toolbox will allow for the greater utilisation of more sophisticated climate analyses within the general scientific community and consequently improve our understanding of the impacts of climate.

## Supporting Information

**S1 File. Simulated data used to test misclassification rates.**Simulated data used to generate Figures 5 - 7.

**S2 File. Simulated data used to test optimum cross-validation folds.** Simulated data used to generate Figure 8.

## Acknowledgments

We are grateful to Nina McLean, Laurie Rijsdijk, Callum Lawson and Lyanne Brouwer for their feedback and help with testing the code.

